# Biochemical Regulation of Brain Kynurenic Acid Synthesis and Inhibition in Rats is Sensitive to the Time of Day

**DOI:** 10.64898/2025.12.01.691600

**Authors:** Courtney J. Wright, Silas A. Buck, Snezana Milosavljevic, Ashley M. Lewis, Nathan T.J. Wagner, Ana Pocivavsek

## Abstract

Neurochemical imbalances, including elevations of the tryptophan metabolite kynurenic acid (KYNA), an endogenous antagonist of glutamatergic and cholinergic receptors, are linked to cognitive and sleep disturbances in psychiatric and neurocognitive disorders. Therapeutic strategies to reduce brain KYNA by inhibiting kynurenine aminotransferase II (KAT II) are under investigation. However, few studies consider time as a biological variable, despite recent evidence that the time of day can affect brain metabolism and drug effectiveness. Therefore, we explore the hypothesis that KYNA formation and synthesis inhibition change throughout the day. Using rats of both sexes, we measured basal KYNA levels and the effects of kynurenine (100 mg/kg, i.p.), to stimulate *de novo* KYNA, and/or PF-04859989 (KAT II inhibitor, 30 mg/kg, s.c.), at the beginning of light or dark phases. Microdialysis was used to assess extracellular KYNA in the dorsal hippocampus, and *ex vivo* assays evaluated KAT enzyme activity in separate animals. Additionally, we examined KYNA levels and the effect of PF-04859989 during acute sleep deprivation in male rats. Regardless of phase, PF-04859989 reduced basal KYNA levels in male but not female rats, yet it reduced kynurenine-stimulated KYNA synthesis in both sexes, demonstrating a context-specific action in female rats. Importantly, we observed a novel effect of phase in males, as kynurenine-induced KYNA synthesis and its inhibition by PF-04859989 were greater during the dark phase than during the light phase. *Ex vivo*, male KAT II activity was higher, and PF-04859989 was more effective, in the dark than in the light phase, suggesting that properties of the KAT II enzyme itself fluctuate with time of day. Finally, sleep deprivation increased extracellular KYNA levels in the light phase, and PF-04859989 fully ameliorated this increase. Overall, our findings highlight the need to consider time-dependent factors when developing therapies impacting KYNA synthesis.

**Lay Summary:** Changes in brain neurochemistry, including elevations in the tryptophan metabolite kynurenic acid (KYNA), are common in brain disorders that present with sleep disturbances and cognitive deficits as symptoms. KYNA interferes with neurotransmission critical for cognition and sleep, so therapeutic strategies to reduce brain KYNA are being pursued. As recent literature has highlighted the impact of time on brain metabolism and drug efficacy, we, for the first time, explored the hypothesis that KYNA formation and synthesis inhibition change throughout the day. We measured extracellular KYNA levels in the brains of male and female rats and stimulated KYNA synthesis with physiological challenges (exogenous kynurenine, the KYNA bioprecursor, or acute sleep deprivation) and/or inhibited KYNA synthesis pharmacologically (PF-04859989, an inhibitor of the KYNA-synthesizing enzyme kynurenine aminotransferase II (KAT II)). KYNA formation, KAT II enzyme activity, and the effectiveness of PF-04859989 varied throughout the day, highlighting time as a key factor modulating KYNA brain metabolism. PF-04859989 reduced KYNA levels under most conditions. Our findings suggest that the timing of KYNA-targeted treatments should be carefully considered in therapeutic strategies to improve cognition and sleep in brain disorders.

## Introduction

Neurochemical imbalances are regularly observed in individuals with neuropsychiatric illnesses, neurocognitive disorders, and systemic infections, and are causally linked to pervasive symptoms like sleep disturbances and cognitive dysfunction in these conditions. Yet, current treatments do not effectively alleviate such symptoms for patients. Sleep deprivation in healthy individuals alters neurochemistry and induces cognitive impairments. Sleep deprivation can disturb behavior and mood, dysregulate the immune system, and increase cardiometabolic risk, all symptoms frequently reported in the neurological conditions above. Improved sleep quality can often reduce these symptoms. Therefore, we posit that understanding mechanisms underlying disrupted neurochemistry and cognitive dysfunction in neurological disease or following sleep deprivation will provide critical insight into developing new and effective therapeutics to address unresolved symptoms in patients.

Brain elevations in kynurenic acid (KYNA), a tryptophan metabolite of the kynurenine pathway (**Figure 1**), are observed in psychiatric patients with schizophrenia (SZ) and bipolar disorder (BPD),^1–5^ neurology patients with neurocognitive dysfunction,^6–11^ and individuals with acute inflammation in the central nervous system.^12–15^ Mechanistic clinical and preclinical studies show that central KYNA levels increase in response to sleep deprivation^16, 17^, psychological stress,^18, 19^ and infection.^20–22^ KYNA acts as an endogenous antagonist of the α7 nicotinic acetylcholine (α7nACh) and N-methyl-d-aspartate (NMDA) glutamate receptors, thereby impacting neurochemical systems that play major roles in modulating cognitive processes and sleep homeostasis.^23, 24^ Elevated KYNA levels in key brain regions, such as the hippocampus, which is critical for learning and sleep-dependent memory consolidation, may be causally related to cognitive and sleep disturbances in these conditions.^25–27^ Preclinical studies support this association, as elevations in KYNA disrupt sleep architecture and hippocampal-dependent cognition, whereas reductions in KYNA improve these behaviors in rats.^28–32^

**Figure 1:**
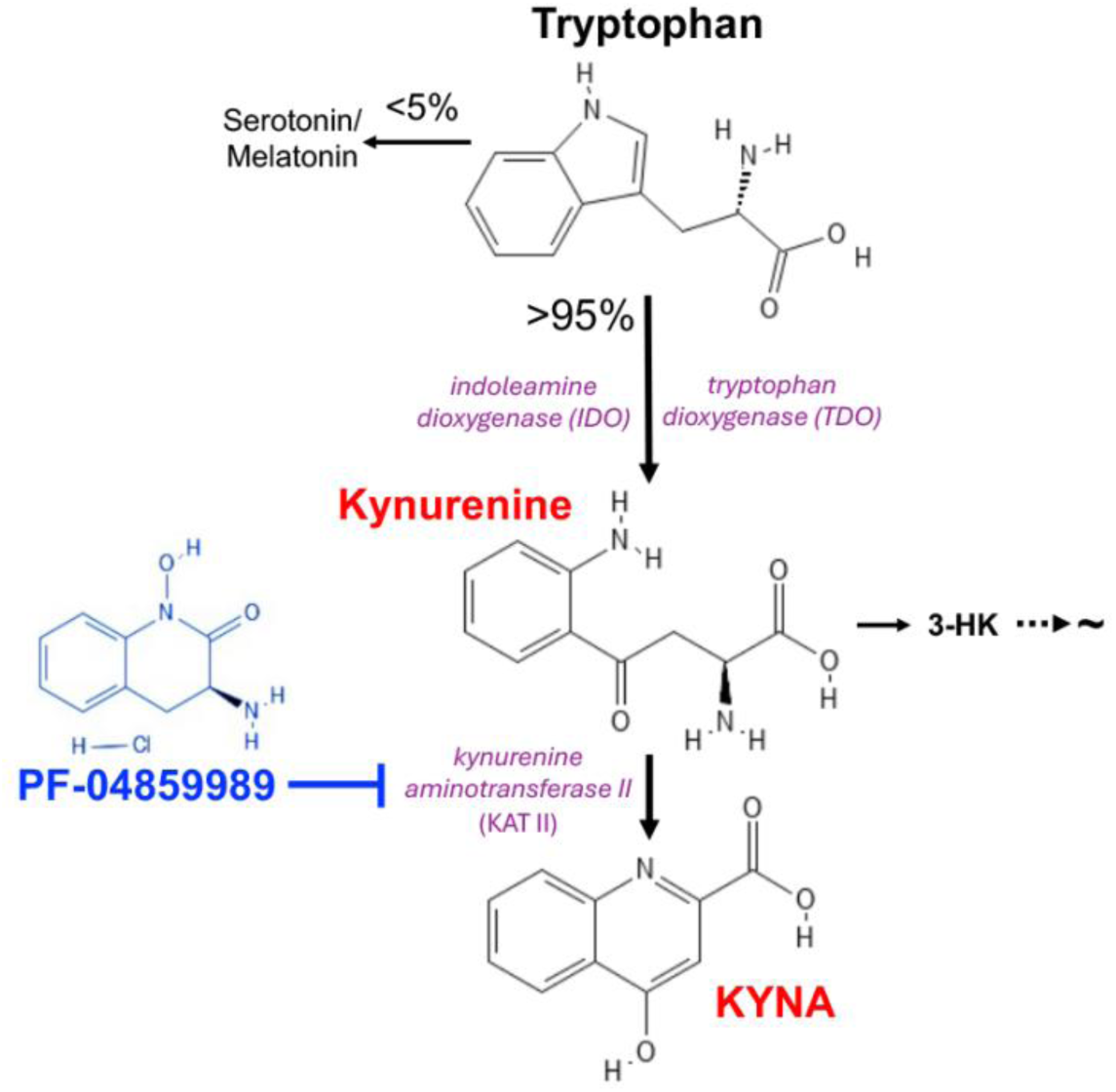
Schematic representation of the kynurenine pathway (KP) of tryptophan degradation, whereby tryptophan is degraded to kynurenine by tryptophan dioxygenase (TDO) or indoleamine dioxygenase (IDO). Subsequently, kynurenine is degraded to 3-hydroxykynurenine (3-HK) or kynurenic acid (KYNA) by kynurenine 3-monooxygenase (KMO) or kynurenine aminotransferase II (KAT II), respectively. PF-04859989 (blue) is an irreversible KAT II inhibitor.

Treatment strategies to reduce brain KYNA levels are being pursued therapeutically (clinicaltrials.gov NCT0401355, NCT06225115).^33, 34^ Translational studies have utilized pharmacological tools that inhibit the main KYNA-synthesizing enzyme in the brain, kynurenine aminotransferase II (KAT II), to successfully reduce brain KYNA, enhance cognitive function, and promote sleep behavior both at baseline and following physiological challenges.^31–33, 35–41^ Yet, our understanding of the influence of circadian phase on neuropharmacology as it relates to modulators of the kynurenine pathway remains limited.

A critical limitation in many studies is the omission of ‘time’ as a variable in the study design, despite recent literature highlighting the significant impact of time of day on brain metabolism and drug efficacy.^42, 43^ Clinical work has shown conspicuous time-of-day differences in peripheral tryptophan metabolism and kynurenine pathway metabolite levels,^44–52^ yet no studies have closely evaluated changes in the brain across 24 hours. We currently use a rat model to fill gaps in our understanding of brain-specific kynurenine pathway metabolism and KYNA synthesis across the day.

Rodents experience ‘active’ and ‘rest’ periods throughout the day. In laboratory settings, when the time of day can be manipulated by controlling the timing of lights on and off, rodent studies are conducted under 12 h light cycles defined by Zeitgeber time (ZT). ZT 0 corresponds to lights on and ZT 12 to lights off. As nocturnal species, rodents spend most of their time asleep when the lights are on (ZT 0-12, “light phase”) and most of their time awake when the lights are off (ZT 12-24, “dark phase”). We have shown that KYNA levels and KYNA-synthesizing enzyme activities in male and female rats change across the light and dark phases.^53, 54^ It is, however, unclear whether central nervous system KYNA synthesis and pharmacological KAT II inhibition are affected by time of day, or how physiological challenges, such as enhanced availability of the bioprecursor kynurenine or acute sleep deprivation (SleepDep), affect this system. Understanding whether time of day affects brain KYNA synthesis can enhance our ability to develop chronotherapeutic approaches, and our present study provides the first steps in this direction. We presently determined *in vivo* and *ex vivo* changes in basal KYNA synthesis across the light and dark cycles and investigated the efficacy of KAT II inhibition with the irreversible, brain-penetrable, and selective (IC_50_ = 230 nM^55^) KAT II inhibitor PF-04859989^35, 36^ in both sexes of rats. Lastly, we explored KYNA synthesis after homeostatic challenge with SleepDep at the beginning of the light phase, demonstrating a rapid increase in extracellular KYNA that PF-04859989 inhibits under these conditions. Taken together, our study highlights ways that KYNA synthesis and inhibition are sensitive to the time of day.

## Results

Male and female rats did not have differences in basal extracellular KYNA levels in the dorsal hippocampus across the light and dark phases. Basal extracellular KYNA levels are reported in **Supplemental Table 1**. A supplemental statistical file reports all statistical results.

### In males, the time of day influences both the effects of KAT II inhibition by PF-04859989 and the synthesis of brain KYNA from exogenous kynurenine

Since previous *in vivo* microdialysis studies on kynurenine pathway neurochemistry have been conducted primarily in male rats, we first examined the time-dependent *de novo* synthesis of KYNA in the dorsal hippocampus of male rats. Peripheral kynurenine challenge (100 mg/kg, to stimulate *de novo* KYNA synthesis in the brain) was administered at either ZT 0, ZT 6, ZT 12, or ZT 18, and extracellular KYNA levels were measured in the dorsal hippocampus. Time of injection significantly influenced the amount of KYNA produced (*F*_3, 23_ = 4.503, *p* < 0.05), and we determined that kynurenine challenge at ZT 12, the beginning of the dark phase, produced twice as much KYNA in the hippocampus as peripheral kynurenine treatment at ZT 0 (*p* < 0.01), the start of the light phase (**Supplemental Figure 1**). We therefore conducted subsequent microdialysis studies to compare treatments at ZT 0 to ZT 12. Furthermore, as our previous studies have noted sex-specific findings in brain KYNA,^16, 54^ we designed the present study to analyze the data separately by sex.

To compare the effects of pharmacologically manipulating KYNA synthesis, either through a kynurenine challenge (as above) or KAT II inhibition using the systemically active, brain penetrant, and irreversible KAT II inhibitor PF-04859989 (30 mg/kg; dose selected based on prior publications and pilot studies in male and female rats^31, 32, 35, 36^), at the start of the light phase (ZT 0) or dark phase (ZT 12), we included the following treatment groups: i) vehicle, ii) PF-04859989, iii) kynurenine, iv) PF-04859989 + kynurenine. Rats received PF-04859989 30 min prior to a kynurenine challenge at ZT 0 or ZT 12 (**Figure 2A**). We first assessed the impact of time on PF-04859989 efficacy of PF-04859989 by comparing PF-04859989 with vehicle. Time of day (“ZT”; light phase: *F*_2.779, 39.18_ = 3.532, *p* < 0.05), PF-04859989 treatment (light phase: *F*_1, 16_ = 4.494, *p* = 0.05; dark phase: *F*_1, 15_ = 4.735, *p* < 0.05) and the interaction between time (ZT) and PF-04859989 treatment (dark phase: *F*_1.953, 24.41_ = 4.772, *p* < 0.05) significantly influenced extracellular KYNA levels in the male dorsal hippocampus (**Figure 2B**). PF-04859989 treatment, irrespective of phase, consistently reduced KYNA levels over time compared to vehicle. Of note, KYNA levels decreased across the light phase and increased across the dark phase under vehicle conditions. Area under the curve (AUC) analysis comparing light and dark phases revealed significant effects of PF-04859989 treatment (*F*_1, 31_ = 9.434, *p* < 0.01; light phase post-hoc: *p* = 0.0817, dark phase post-hoc: *p* < 0.05) (**Figure 2C**). PF-04859989 reduced KYNA by 36-48%.

**Figure 2:**
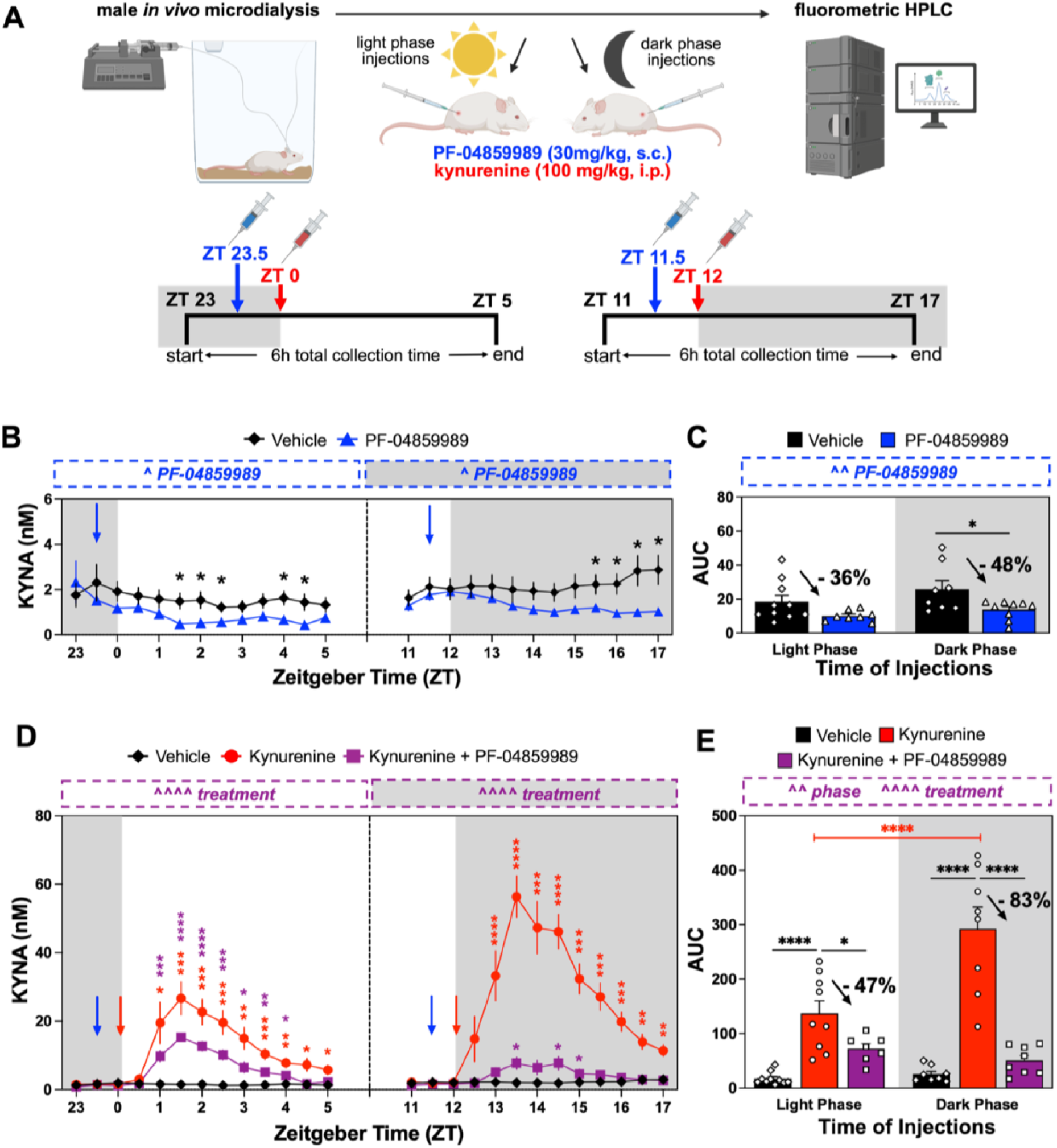
In male rats, compared to light phase, dark phase kynurenine challenge (100 mg/kg, i.p) stimulates more extracellular kynurenic acid (KYNA) in the hippocampus, and pre-treatment with PF-04859989 (30 mg/kg, s.c.) inhibits KYNA levels to a greater extent. **(A)** Experimental timeline. Male rats were peripherally injected with kynurenine to stimulate *de novo* KYNA at the beginning of the light phase (Zeitgeber time (ZT) 0) or dark phase (ZT 12). Injection times are denoted by red arrows. PF-04859989, a systemically active brain-penetrant kynurenine aminotransferase (KAT II) inhibitor, was administered to inhibit KYNA synthesis 30 min prior (ZT 23.5 or ZT 11.5). Injection times are denoted by blue arrows. Extracellular KYNA collected via *in vivo* microdialysis in the dorsal hippocampus was evaluated in 30-minute fractions for the duration of the experiment. **(B)** Extracellular KYNA following vehicle or PF-04859989 treatment. **(C)** Area under the curve (AUC) of extracellular KYNA following vehicle or PF-04859989 treatment. **(D)** Extracellular KYNA following kynurenine challenge ± PF-04859989 pretreatment. **(E)** AUC of extracellular KYNA following kynurenine challenge ± PF-04859989 pretreatment. All data are mean ± SEM. Two-way RM ANOVA separated by phase (panels B/D) or two-way ANOVA (panels C/E). Significant main effects of PF-04859989, treatment, and/or light phase are displayed in boxed annotations above each figure panel: ^ *p* < 0.05, ^^ *p* < 0.01, ^^^^ *p* < 0.0001; Fisher’s LSD post hoc test (panels B/C/E) or Dunnett’s post hoc test to vehicle (panel D): \**p* < 0.05, \*\**p* < 0.01, *** *p* < 0.001, \*\*\*\**p* < 0.0001. N = 7 - 10 per group.

Next, we assessed whether KYNA synthesis from exogenous kynurenine (kynurenine challenge) and the effects of PF-04859989 pretreatment on this response are influenced by time of day. Kynurenine challenge transiently elevated KYNA levels in both the light and dark phases, and pretreatment with PF-04859989 significantly blunted this response (**Figure 2D**). When assessing each phase independently and comparing treatment groups across time (ZT), we determined significant ZT x treatment interactions (light phase: *F*_4.174, 45.91_ = 10.06, *p* < 0.0001; dark phase: *F*_6.680, 68.75_ = 22.23, *p* < 0.0001), as well as main effects of ZT (light phase: *F*_2.087, 45.91_ = 26.29, *p* < 0.0001; dark phase: *F*_2, 23_ = 51.01, *p* < 0.0001) and treatment (light phase: *F*_2, 24_ = 17.22, *p* < 0.0001; dark phase: *F*_3.340, 68.75_ = 30.96, *p* < 0.0001). AUC analysis revealed a significant phase x treatment interaction (*F*_2, 44_ = 11.68, *p* < 0.0001) and main effects of both phase (*F*_1, 44_ = 8.761, *p* < 0.001) and treatment (*F*_2, 44_ = 56.46, *p <* 0.0001) on KYNA levels (**Figure 2E**). Kynurenine challenge significantly elevated KYNA levels by 8-fold (*p* < 0.0001) during the light phase and 11.4-fold (*p* < 0.0001) during the dark phase relative to vehicle treatment. Notably, kynurenine challenge during the dark phase produced significantly greater *de novo* extracellular KYNA than during the light phase (*p* < 0.0001). Regardless of phase, PF-04859989 pretreatment significantly attenuated KYNA levels compared to kynurenine alone (light phase: *p* < 0.05, dark phase: *p* < 0.0001). However, when PF-04859989 was given prior to the dark phase, the reduction in *de novo* KYNA production following kynurenine challenge was more pronounced than when the KAT II inhibitor was given before the light phase.

### In females, the ability of the KAT II inhibitor PF-04859989 to block kynurenine-induced KYNA synthesis in the brain is significantly influenced by the time of day

To expand our understanding of the impact of biological sex on KP metabolism, we conducted a microdialysis study in female rats (**Figure 3A**). To our knowledge, no prior studies have evaluated whether PF-04859989 can reduce basal extracellular KYNA levels in female rats. Surprisingly, our results demonstrate that the current dose of PF-04859989 fails to reduce basal KYNA levels in the dorsal hippocampus of female rats in either the light or dark phases (**Figure 3B**). Vehicle-treated or PF-04859989-treated females showed no differences in KYNA levels between the light and dark phases in our AUC analysis (**Figure 3C**). Additionally, KYNA levels in females were not significantly influenced by biological sex (**Supplemental Table 1**) or the estrous cycle (**Supplemental Figure 2**).

**Figure 3:**
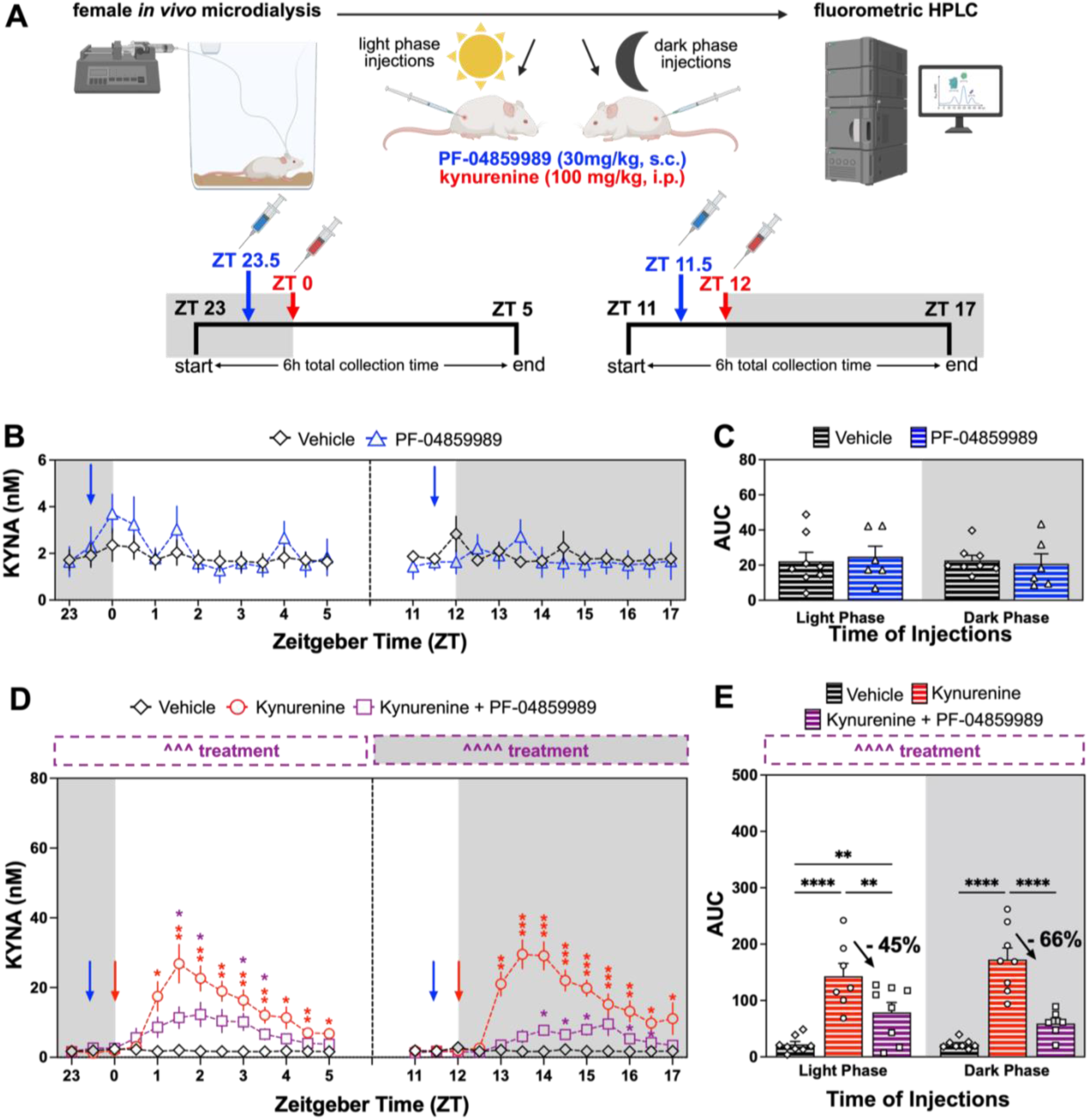
In female rats, light and dark phase kynurenine challenge (100 mg/kg, i.p.) equivocally stimulates extracellular kynurenic acid (KYNA) in the hippocampus, yet PF-04859989 pretreatment (30 mg/kg, s.c.) inhibits KYNA levels to a greater extent in the dark phase. **(A)** Experimental timeline. Female rats were peripherally injected with kynurenine to stimulate *de novo* KYNA at the beginning of the light phase (Zeitgeber time (ZT) 0) or dark phase (ZT 12). Injection times are denoted by red arrows. PF-04859989, a systemically active brain-penetrant kynurenine aminotransferase (KAT II) inhibitor, was administered to inhibit KYNA synthesis 30 min prior (ZT 23.5 or ZT 11.5). Injection times are denoted by blue arrows. Extracellular KYNA collected via *in vivo* microdialysis in the dorsal hippocampus was evaluated in 30-minute fractions for the duration of the experiment. **(B)** Extracellular KYNA following vehicle or PF-04859989 treatment. **(C)** Area under the curve (AUC) of extracellular KYNA following vehicle or PF-04859989 treatment. **(D)** Extracellular KYNA following kynurenine challenge ± PF-04859989 pretreatment. **(E)** AUC of extracellular KYNA following kynurenine challenge ± PF-04859989 pretreatment. All data are mean ± SEM. Two-way RM ANOVA separated by phase (panels B/D) or two-way ANOVA (panels C/E). Significant main effects of PF-04859989, treatment, and/or light phase are displayed in boxed annotations above each figure panel: ^^^ *p* < 0.001, ^^^^ *p* < 0.0001. Fisher’s LSD post hoc test (panels B/C/E) or Dunnett’s post hoc test to vehicle (panel D): \**p* < 0.05, \*\**p* < 0.01, *** *p* < 0.001, \*\*\*\**p* < 0.0001. N = 6 – 9 per group.

In females, kynurenine challenge transiently elevated KYNA levels, and pretreatment with PF-04859989 blunted this response regardless of phase (**Figure 3D**). KYNA levels were influenced by interactions between time (ZT) and treatment (light phase: *F*_5.212, 50.17_ = 8.490, *p* < 0.0001; dark phase: *F*_5.889, 55.45_ = 12.65, *p* < 0.0001), as well as main effects of ZT (light phase: *F_2_*_.606, 50.17_ = 21.31, *p* < 0.0001; dark phase: *F*_2.944, 55.45_ = 19.55, *p* < 0.0001) and treatment (light phase: *F*_2, 20_ = 12.48, *p* < 0.001; dark phase: *F*_2, 19_ = 31.30, *p* < 0.0001). AUC analysis revealed that treatment (*F*_2, 39_ = 42.09, *p* < 0.0001), but not phase, affected kynurenine-induced KYNA synthesis and its inhibition by PF-04859989 in females (**Figure 3E**). Kynurenine challenge significantly increased KYNA 6.5-fold during the light phase (*p* < 0.0001) and 7.8-fold during the dark phase (*p* < 0.0001) compared to vehicle treatment. PF-04859989 pretreatment was more effective at preventing kynurenine-induced KYNA synthesis in the dark phase (PF-04859989 + kynurenine vs vehicle: not significant, *p* > 0.05) than in the light phase (PF-04859989 + kynurenine vs vehicle: *p* < 0.01).

### Brain KAT II enzyme activity exhibits diurnal variation

We next sought to determine whether time-dependent alterations in KAT II activity underlie the previously observed differences in extracellular KYNA. To this end we conducted *ex vivo* KAT enzyme activity assays on male and female forebrain homogenates which included the whole hippocampus region collected at 3-hour intervals for 24 hours (**Figure 4A**). We focused on the KAT II isoform, the predominant KYNA-synthesizing enzyme in the brain,^56^ though we also conducted KAT I enzyme activity assays (**Supplemental Figure 3**). ZT significantly affected KAT II enzyme activity in both male (*F*_7, 24_ = 6.428, *p* < 0.001; **Figure 4B**) and female (*F*_7, 24_ = 5.495, *p* < 0.001; **Figure 4C**) samples, demonstrating broad variation in KAT II activity across the day. Similarly, male and female KAT I activity was significantly influenced by ZT (**Supplemental Figure 3**).

**Figure 4:**
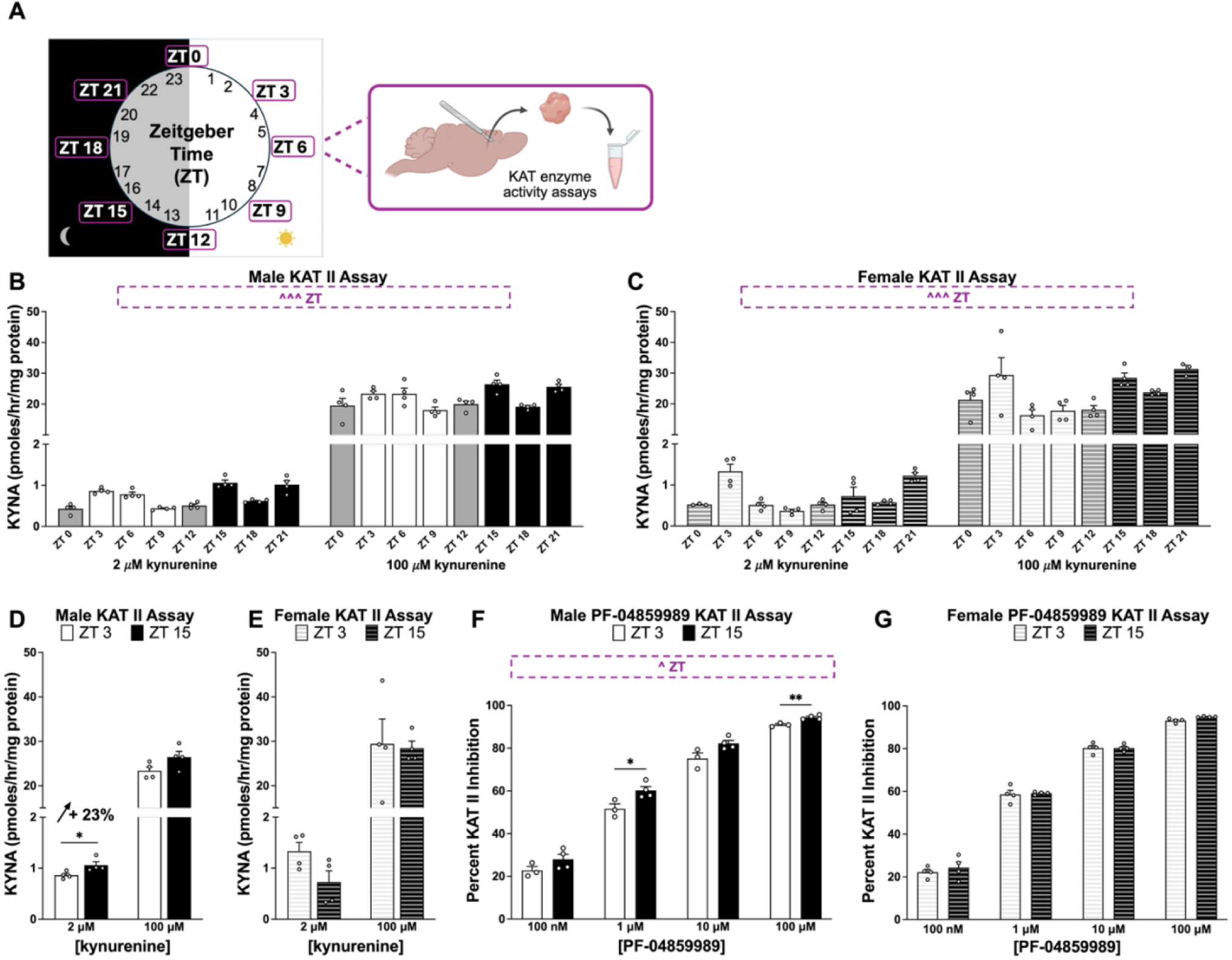
Kynurenine aminotransferase II (KAT II) enzyme activity changes across the time of day. In males, KAT II enzyme activity and its inhibition by PF-04859989 are significantly altered between Zeitgeber time (ZT) 3 and ZT 15. **(A)** Experimental approach. Brain samples were collected from male and female rats at 3-hour intervals for *ex vivo* enzyme activity assays, which were assessed by measuring kynurenic acid (KYNA) production. **(B)** Male forebrain KAT II enzyme activity. **(C)** Female forebrain KAT II enzyme activity. **(D)** Male forebrain KAT II enzyme activity at ZT 3 and ZT 15. **(E)** Female forebrain KAT II enzyme activity at ZT 3 and ZT 15. **(F)** Male percent of KAT II inhibition with PF-04859989 at ZT 3 and ZT 15. **(G)** Female percent of KAT II inhibition with PF-04859989 at ZT 3 and ZT 15. All data are mean ± SEM. Two-way RM ANOVA (panels B/C/F/G). Significant main effects of time (“ZT”) are displayed in boxed annotations above each figure panel: ^ p < 0.05, ^^^ p < 0.001. Mann-Whitney tests (panels D/E) or Fisher’s LSD post-hoc (panels F/G): \**p* < 0.05, \*\**p* < 0.01. N = 3 – 4 per group.

We next compared KAT II activity at ZT 3 and ZT 15, the first tissues collected after the light and dark phase transitions, respectively, separately by sex. In males, KAT II enzyme activity at ZT 15 was 23% higher than at ZT 3 when assayed with 2 μM kynurenine (**Figure 4D**, *p* < 0.05). In contrast, no difference in KAT II activity was observed between these time points in females (**Figure 4E**). KAT I activity was reduced in females but not males at ZT 15 compared to ZT 3 (**Supplemental Figure 3**).

We also assessed the ability of PF-04859989 to inhibit *ex vivo* KAT II activity at these time points in a dose-response manner. At all tested concentrations, PF-04859989 significantly reduced KAT II enzyme activity compared to the active control in males and females (**Supplemental Figure 4**). In males, both concentration (*F*_1.690, 8.451_ = 831.9, *p* < 0.0001) and ZT (*F*_1, 5_ = 10.62, *p* < 0.05) significantly influenced the percent inhibition of KAT II activity by PF-04859989 (**Figure 4F**). At 1 μM and 100 μM concentrations, PF-04859989 produced significantly greater inhibition of KAT II activity at ZT 15 compared to ZT 3 (1μM: *p* < 0.05, 100 μM: *p* < 0.01). In females, no differences in inhibition were observed between ZT 3 and ZT 15, although the expected concentration-dependent effect of PF-04859989 (*F*_1.822, 10.93_ = 1176, *p* < 0.0001) was evident (**Figure 4G**).

### PF-04859989, KAT II inhibitor, prevents elevated extracellular KYNA induced by acute sleep deprivation (SleepDep) in males

In addition to exogenous kynurenine, we investigated whether KAT II inhibition could prevent KYNA elevations induced by acute SleepDep via gentle handling which is a physiological challenge previously shown to increase brain KYNA in male but not female rats.^16^ Thereby, we investigated whether PF-04859989 (30 mg/kg) could prevent SleepDep-induced elevations in KYNA levels in males (**Figure 5A**). SleepDep was restricted to the beginning of the light phase (ZT 0 - 6), when rats typically obtain most of their sleep, and sleep pressure is highest. Monitoring of extracellular KYNA was extended through ZT 12 to capture recovery from SleepDep.

**Figure 5:**
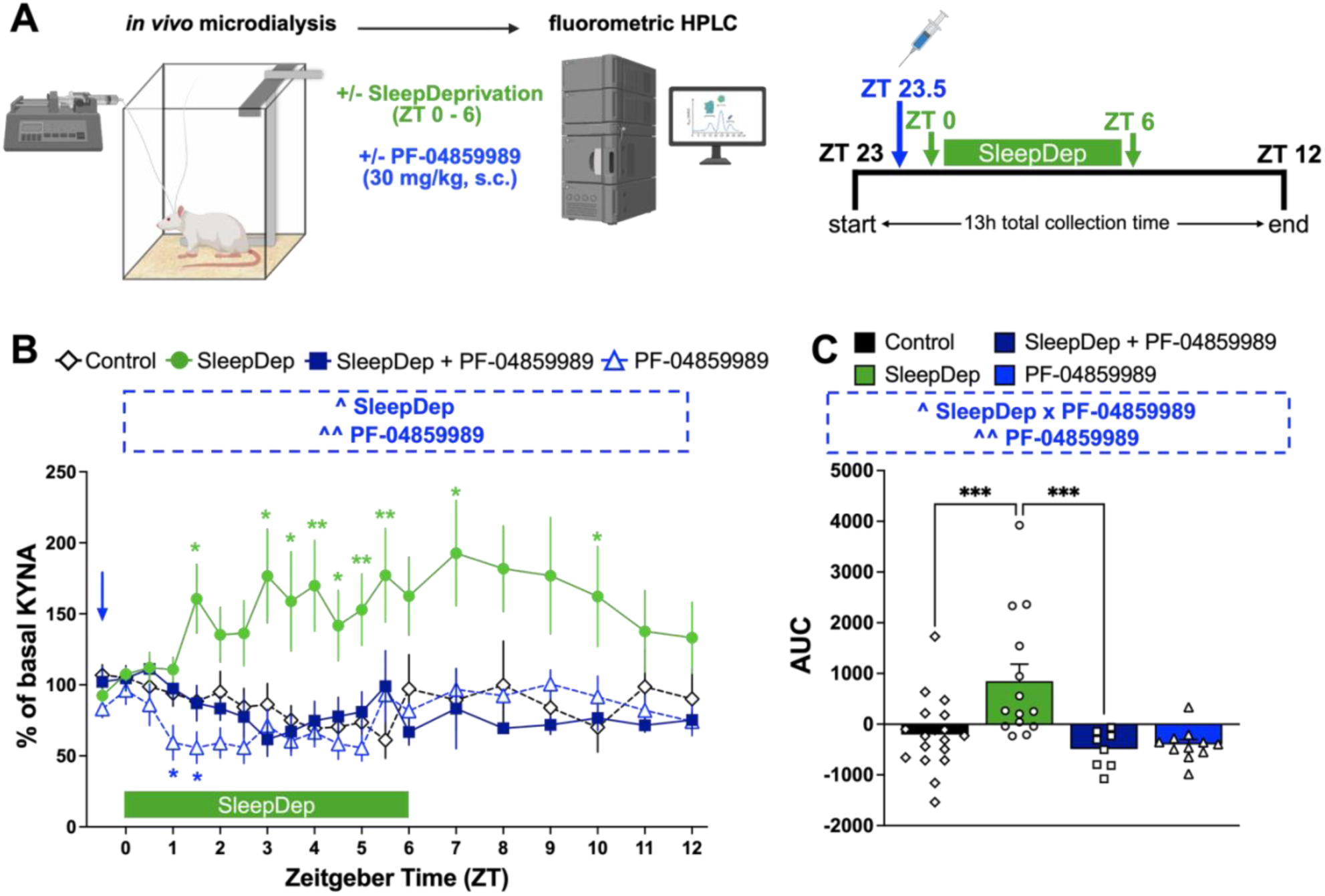
Accumulation of extracellular KYNA with sleep deprivation (SleepDep) is prevented with PF-04859989 pretreatment in males. **(A)** Experimental timeline. Male rats were sleep-deprived from Zeitgeber time (ZT) 0 to ZT 6, as denoted by the green arrows, using a manual, gentle bar-sweeping approach. PF-04859989 (30 mg/kg, s.c.), a systemically active kynurenine aminotransferase II (KAT II) inhibitor, was administered 30 min before SleepDep (ZT 23.5) to prevent KYNA synthesis. Injection time is denoted by a blue arrow. For the duration of the experiment, microdialysis was performed in the dorsal hippocampus to assess extracellular KYNA levels. **(B)** Extracellular KYNA levels are shown as a percent change from baseline following control, PF-04859989, SleepDep, and SleepDep with PF-04859989. **(C)** Area under the curve (AUC) of extracellular KYNA. Three-way ANOVA (panel B) or two-way ANOVA (panel C). Significant interactions and main effects of PF-04859989 and/or SleepDep are displayed in boxed annotations above each figure panel: ^ p < 0.05, ^^ p < 0.01. Fisher’s LSD post-hoc test: \**p* < 0.05, \*\**p* < 0.01, \*\*\**p* < 0.001. N = 8 – 17 per group.

Our analysis revealed that SleepDep (*F*_1, 47_ = 6.894, *p* < 0.05) and PF-04859989 (*F*_1, 47_ = 8.455, *p* < 0.01) significantly affected extracellular KYNA levels in the hippocampus (**Figure 5B**). Acute SleepDep elevated extracellular KYNA levels throughout most of the light phase compared with control, whereas PF-04859989 treatment prevented this increase. KYNA levels of sleep-deprived males treated with PF-04859989 were not significantly different from those of control (*ad libitum* sleep) vehicle-treated or control PF-04859989-treated males. With AUC analysis, an interaction between SleepDep and PF-04859989 (F_1, 46_ = 5.632, *p* < 0.05) and a main effect of PF-04859989 (F_1, 46_ = 10.15, *p* < 0.01) influenced KYNA levels (**Figure 5C**). Our post-hoc analysis confirmed that SleepDep elevated KYNA levels (control vs SleepDep: *p* < 0.001) and that pretreatment with PF-04859989 effectively blocked this increase (PF-04859989 + SleepDep vs SleepDep: *p* < 0.001).

## Discussion

Our current study found that time of day significantly affected *de novo* KYNA production and the inhibition of KYNA synthesis by PF-04859989 in the rat brain. These time-of-day effects were more pronounced in males than in females; however, in both sexes, the activity of the KYNA-synthesizing enzymes KAT I and KAT II varied across the day. PF-04859989 treatment could ameliorate SleepDep-induced elevations of KYNA in male rats.

Extracellular hippocampal KYNA levels decreased across the light phase and increased across the dark phase in male rats, consistent with previously reported literature indicating that tryptophan and its metabolites exhibit circadian fluctuations both centrally and peripherally.^44–49, 51, 53, 54^ Both tryptophan and kynurenine can cross the blood-brain barrier (BBB), and peripheral levels readily influence brain kynurenine pathway metabolism.^9^ As an essential amino acid obtained through food, peripheral tryptophan levels are greater in the dark phase than in the light phase in rodents.^57^ Thereby, the *ad libitum* access to food in the present study may have influenced circulating kynurenine levels; however, our experimental approach relied on stimulating KYNA levels via exogenous peripheral kynurenine administration and allowed us to consider brain-specific mechanisms underlying the diurnal changes in hippocampal extracellular KYNA levels.

The phase-dependent changes in basal extracellular KYNA we observed could be due to differences in efflux or transporter activity between the light and dark phases, though these hypotheses remain to be tested experimentally. Efflux of KYNA from the brain occurs via organic anion transporters, an active transport mechanism,^27, 58^ and studies suggest enhanced brain efflux during sleep.^59, 60^ Clearance mechanisms during sleep may thereby contribute to the stable extracellular KYNA levels that we observed across the light phase when animals were left undisturbed. In contrast, reduced brain efflux during waking may contribute to the increased KYNA levels we observed across the dark phase, and to an even greater extent in SleepDep experiments, as KYNA appears to accumulate in the extracellular space. Given increasing evidence for sex differences in sleep regulation and current results showing that males, but not females, accumulate KYNA with sustained wakefulness,^16^ investigating what clearance mechanisms drive sex-dependent responses to SleepDep is an important, and translationally relevant, future direction. As KYNA does not readily cross the BBB and is primarily synthesized from plasma kynurenine, which enters the brain via the large neutral amino acid transporter,^61, 62^ circadian changes in transporter function or BBB permeability could influence brain substrate availability and, therefore, possibly KYNA metabolism.^63–65^ We were unable to detect extracellular kynurenine in our samples, nor did we measure plasma kynurenine, so further investigation is warranted to directly test this hypothesis. Additionally, regulation of kynurenine and KYNA transfer into or out of the astrocytes via sodium-independent processes or organic anion transporters,^58, 61^ respectively, could also contribute to these phase-dependent changes.

It is important to note that our current findings are based on extracellular microdialysis measurements in the dorsal hippocampus; KYNA levels in tissue homogenates or different brain regions may exhibit slightly different dynamics across the circadian cycle, reflecting distinct pools and regulatory mechanisms between intracellular and extracellular compartments ^58^ and unique microenvironments between brain regions.^66–68^ Variable circadian influences across brain regions have been observed,^69, 70^ and our study in the hippocampus provides the first evidence of these dynamics with regards to KYNA synthesis. Despite its ubiquitous presence as the primary synthesizing enzyme for cerebral KYNA production, KAT II is not equally expressed throughout the brain. It is highly localized to specific regions, particularly in astrocytes within germinal zone and peripheral kynurenine challenge produces differential amounts of KYNA across regions.^32, 56, 71, 72^ The hippocampus consistently produces high *de novo* KYNA and combined with behavioral studies, ^29, 39, 73^ these data suggest a strong, and potentially unique, role for KYNA in this region.

In our current study, we observed variation in *ex vivo* KAT I and KAT II enzyme activity across the light and dark phases. *In vivo* findings showed a marked increase in KYNA formation during the dark phase compared to the light phase in males, whereas females did not exhibit this phase-dependent difference. *Ex vivo* assays revealed greater KAT II activity at ZT 15 than at ZT 3 in male rat brains. In females, KAT II activity did not differ substantially across these time points, yet KAT I activity was reduced at ZT 15 compared with ZT 3. Because KAT II contributes approximately 60% of *de novo* KYNA synthesis,^56, 74^ we interpret that increased KAT II activity during the dark phase drives the elevated KYNA formation observed in males. The *ex vivo* assays were conducted in forebrain homogenates that included the entire hippocampus, allowing KAT I and II assays in the same sample but preventing us from using only the hippocampus, a minor limitation for understanding diurnal KAT activity in this brain region. The distinct biochemical properties of KAT I and KAT II, including differences in substrate preferences, pH optimum, utilization of α-keto acids, competing endogenous substrates, and enzymatic functions, may underlie their divergent circadian patterns.^27^ Additionally, regulators of kynurenine pathway enzymes, such as steroids, growth factors, and immune signaling molecules, fluctuate with time of day^75, 76^ and may further modulate brain kynurenine metabolism, though this remains to be tested in the brain.^77, 78^ While our study focused on the direct KYNA synthesizers, KATs, information currently remains limited about the circadian regulation of the alternative branch of kynurenine metabolism, initiated by kynurenine 3-monooxygenase (KMO). It is feasible for KMO to indirectly influence central KYNA synthesis by affecting kynurenine flux,^79, 80^ and further understanding of circadian influences on this mechanism poses important future questions.

Time-dependent changes in cellular conditions may further contribute to the circadian pattern of KYNA synthesis. Variations in glucose availability, oxidative stress, and reactive oxygen species (ROS) production across the light-dark cycle could influence both enzymatic and non-enzymatic KYNA formation.^81^ Enzymatic KYNA metabolism is highly dependent on glucose availability,^82^ which varies as a function of sleep-wake states and, therefore, ZT.^81, 83^ Kynurenine can also be converted to KYNA through non-enzymatic oxidative pathways mediated by ROS.^84^ These reactive species can accumulate during sleep deprivation^85^ and are known to reach peak levels during the dark phase,^86^ coinciding with the period of elevated KYNA levels observed in our study. Thus, time-dependent glucose availability and ROS-mediated KYNA metabolism are important avenues for future experiments.

When we inhibited KAT II with PF-04859989, we observed a more pronounced blockade of *de novo* KYNA production after kynurenine challenge in the dark than in the light phase, suggesting that the drug or enzyme itself exhibits time-of-day-dependent properties. PF-04859989 acts by covalently binding the essential pyridoxal 5’-phosphate (P5P or PLP) cofactor in the KAT II active site. Thus, theoretically, the *in vivo* findings could be attributed to increased brain drug transfer or cofactor availability during the dark phase.^60, 87^ Without pharmacokinetic and pharmacodynamic profiling, including measurements of plasma and brain concentrations over time, this remains to be explicitly determined. Yet *ex vivo* enzyme assays, in which drug and cofactor concentrations are controlled, also supported the notion that PF-04859989 inhibition of KYNA synthesis was greater in tissue collected during the dark phase than during the light phase. Therefore, we speculate that the drug’s ability to bind P5P may differ across time points.^55^ Together, these findings suggest that dosing PF-04859989 before the dark phase may be most effective in reducing brain KYNA levels, yet further biochemical and functional studies are warranted to better understand the behavioral implications and, thereby, any time-of-day differences that may govern specific behaviors, such as sleep or cognition.

Differences in *de novo* KYNA production, particularly the effectiveness of PF-04859989, in female rats deserve further discussion. In females, PF-04859989 administered before a kynurenine challenge significantly reduced KYNA levels across phases; however, it failed to reduce basal extracellular KYNA levels. Therefore, we identify a novel context-specific pharmacological profile of PF-04859989 that may warrant further investigation of pharmacological means to inhibit basal KYNA levels in the female brain. The lack of effect of PF-04859989 basally was not explained by differences in baseline KYNA between sexes, fluctuations in KYNA across estrous phases, or any deficiency in PF-04859989’s ability to inhibit KAT II *ex vivo* in females. Estrogen and its phosphate and sulfur ester forms have been described as reversible inhibitors of KAT II^88, 89^ and may compete with PF-04859989 for access to the enzyme’s active site,^88^ thereby limiting its inhibitory efficacy. It is therefore possible that higher doses of PF-04859989 might overcome this competition in females. PF-04859989 has low reactivity with other KAT isoforms and P5P-dependent enzymes,^36, 55^ yet higher doses could eventually compromise KAT II selectivity and promote off-target effects. More likely, because PF-04859989 covalently binds P5P, increasing PF-04859989 dosage could reduce the activity of enzymes that also rely on P5P as a cofactor.

In addition to direct enzymatic effects, sex-dependent differences in compound and metabolite transport may contribute to these findings.^90^ PF-04859989 crosses the BBB, however, BBB permeability is itself influenced by biological sex and sex hormones.^91, 92^ Estrogen has been shown to decrease BBB permeability in reproductively intact female rats,^93^ which could limit drug penetration into the brain during periods of elevated estrogen. Moreover, the transporters responsible for KYNA movement across cellular membranes, organic anion transporters (OATs),^58^ exhibit sex differences in expression and activity in the kidneys,^94, 95^ wherein renal cortical OAT1 and OAT3 expression is enhanced by androgens and suppressed by estrogens, suggesting a broader hormonal regulation that could extend to the brain and warrants further investigation in regard to the transport of KYNA.^96^ Additionally, although time-of-day-dependent changes in these transports have not yet been described, such regulation could influence the extent to which KYNA is released into or cleared from the extracellular space. Together, these factors, if validated by future studies, may interact to shape the sex-specific and potentially circadian dynamics of KYNA metabolism and pharmacological inhibition.

Interestingly, in females, KYNA synthesis from kynurenine was comparable between the light and dark phases, consistent with the absence of time-of-day differences in *ex vivo* KAT II enzyme activity. In addition to enzymatic conversion by KATs, KYNA can be synthesized through oxidative mechanisms.^84^ Males can express higher ROS and oxidative stress than females, potentially due to estrogen’s antioxidant and neuroprotective roles.^97, 98^ Estrogen can also act as a regulator of circadian rhythmicity by modulating clock gene expression,^99^ which may, in turn, influence KAT II expression and activity in a time-of-day-dependent manner. Together, these factors may contribute to the absence of a clear circadian pattern in KYNA production observed in females and deserve consideration for future studies.

Clinically, elevated KYNA levels,^1–14^ along with sleep disruptions^100–102^ and cognitive impairments,^103–105^ are well-documented in neuropsychiatric and neurocognitive disorders, as well as in chronic infections. Dysfunction of the hippocampus, a brain region crucial for learning, memory, and sleep regulation, may underlie these symptoms. Because elevated KYNA has been implicated in cognitive and sleep abnormalities, ^9, 25–27^our microdialysis studies conducted in the dorsal hippocampus provide translationally relevant insight into mechanisms that may contribute to clinical dysfunction.

Inhibiting KYNA synthesis is a promising therapeutic strategy for mitigating cognitive and sleep disturbances associated with psychiatric disease and sleep deprivation.^33^ Preclinical evidence shows that KAT II inhibitors, including PF-04859989, reduce brain KYNA levels and improve cognition in male rats.^37–39^ PF-04859989 has been shown to prevent stimulant-induced deficits in auditory gating, working memory, spatial memory, and sustained attention^36^; to attenuate KYNA-mediated impairments in fear discrimination after stress exposure^40^; and to facilitate long-term potentiation in mice with elevated KYNA.^41^ Although these findings strongly support the compound’s pro-cognitive potential, prior studies have largely focused on males. Further, considering that both cognitive performance and KYNA metabolism exhibit circadian variation, future behavioral studies of KAT II inhibition should incorporate time of day and biological sex as key experimental variables. Moreover, PF-04859989, administered at the onset of either the light or dark phase, enhances sleep in both sexes,^31^ suggesting that KAT II inhibition may represent an effective chronotherapeutic approach to improve sleep quality.

In summary, our study demonstrates that the effects of PF-04859989 on KYNA synthesis are influenced by both time of day and biological sex. These data highlight the importance of circadian timing and sex-specific factors when evaluating KAT II inhibitors as potential treatments for disorders characterized by dysregulated KYNA metabolism.

## Methods

### Animals

Adult male and female Wistar rats (200 - 300g) were obtained from Charles River Laboratories. Animals received standard rodent chow, Tekland Global 18% protein extruded rodent diet, which contained 0.2% tryptophan (Envigo, Cumberland, Virginia) *ad libitum* and were given at least one week to acclimate to the animal facility before use in any experiment. Animals were kept on a 12/12-hour light/dark cycle, where lights on corresponded to Zeitgeber time (ZT) 0 and lights off to ZT 12. All animals were housed in a temperature-controlled facility fully accredited by the American Association for the Accreditation of Laboratory Animal Care. All protocols were approved by the Institutional Animal Care and Use Committee at the University of South Carolina School of Medicine and were in accordance with the National Institutes of Health Guide for the Care and Use of Laboratory Animals.

### Chemicals

L-Kynurenine sulfate (“kynurenine,” purity: 99.4%) was obtained from Sai Advantium (Hyderabad, India). PF-04959989 was obtained from WuXi AppTec (Shanghai, China). All other chemicals were obtained at the highest commercially available purity from various suppliers. PF-04859989 and kynurenine were prepared for injections daily by dissolving 10 mg/mL in ultrapure water or phosphate-buffered saline, respectively. For kynurenine, the pH was adjusted to 7.2-7.6 with 0.1 N sodium hydroxide. Vehicle injections were ultrapure water (s.c.) or saline (i.p.).

### Experiment 1: PF-04859989 and Kynurenine Challenge Microdialysis

#### Surgery

Animals were placed on a stereotaxic frame (Stoelting Co. Wood Dale IL, USA) under isoflurane anesthesia (2 - 5%) to implant a guide cannula (1.0 mm outer diameter, SciPro Inc., Sanborn, NY, USA) over the dorsal hippocampus (AP: −3.4 mm; LM: ±2.3 mm; DV: −1.5 mm from bregma). Carprofen (5 mg/kg, s.c.) was given for analgesic support. Two surgical screws were inserted into 0.5 mm burr holes on the skull, and acrylic dental cement secured each cannula in place. Animals were given 24-48 hours of recovery before experiments.

#### Intrahippocampal Microdialysis

On the day of experiments, at ZT 22 or ZT 10, a microdialysis probe (2 mm PES membrane/14 mm shaft/6kD, SciPro Inc.) was inserted through the guide cannula and Ringer solution (147 mM NaCl, 4 mM KCl, 1.4 mM CaCl_2_) was perfused at a flow rate of 2.5 μL/minute with a microperfusion pump (Harvard Apparatus, Holliston, MA, USA). After 30 minutes, the flow rate was reduced to 1.0 μL/minute for the remainder of the experiment. Samples were collected in 30-minute fractions in a refrigerated autosampler (CMA Microdialysis AB - Harvard Apparatus Inc., Holliston, MA, USA). After one hour of baseline collection, animals received either PF-04859989 (30 mg/kg, s.c.) or vehicle (s.c.) injections at ZT 23.5 (prior to the start of light phase) or ZT 11.5 (prior to the start of dark phase). Animals then received either kynurenine (100 mg/kg, i.p.) or vehicle (i.p.) injections at ZT 0 or ZT 12. Collection continued in 30-minute fractions for 5 hours post-injection. Food was provided during all microdialysis experiments, though intake was not monitored. Dialysate samples were stored at −80°C until biochemical analyses.

#### Placement Verification

At the end of the microdialysis experiment, brains were removed from euthanized animals and drop-fixed in 10% formalin solution to verify the correct placement of the guide cannula and probe. Brains were moved to 20% sucrose, then 40 μm coronal sections obtained on a cryostat (Microm HM 560, Leica BioSystems, Nussloch, Germany) containing the dorsal hippocampus were collected on gelatin-coated slides. Slides were stained with Neutral Red and evaluated under a light microscope (Laxco, SeBa Pro 4) to confirm probe placement.

### Experiment 2: KAT Enzyme Activity Assays

#### Tissue Collection

Animals were euthanized via CO_2_ asphyxiation every three hours across the 24-hour light/dark cycle, beginning at ZT 0. Brains were promptly removed, snap frozen on dry ice, and stored at −80°C.

#### KAT I/KAT II Assays

In an effort to conduct KAT I and KAT II enzyme assays from the same tissue samples, whole forebrain homogenates which encompassed the entire hippocampal fraction were used for assays. On the day of the assays, forebrains were thawed on wet ice and homogenized (1:5 w/v) in ultrapure water. A portion of the homogenate was diluted 1:10 with homogenization buffer (5 mM Tris acetate buffer pH 8.0 containing 50 μM pyridoxal 5’-phosphate (P5P) and 10 mM β-mercaptoethanol) and dialyzed against the homogenization buffer overnight at 4°C. The remaining 1:5 homogenate was stored at −80°C for later use. The next day, 80 μL of dialyzed homogenate (1:20 final dilution) was combined with 20 μL ultrapure water, and 100 μL reaction cocktail (KAT I: 2 μM or 100 μM kynurenine, 80 μM P5P, 1 mM pyruvate, and 150 mM AMP buffer pH 9.5; KAT II: 2 μM (i.e. << K_m_^106^) or 100 μM kynurenine,^107^ 80 μM P5P,1 mM pyruvate, and 150 mM Tris acetate buffer pH 7.4) and incubated in a 37°C water bath for 2 hours. Blank controls were assessed by adding 20 μL of 1 mM aminooxyacetate, a general KAT inhibitor,^108^ to the reaction instead of water. For assays with the KAT II inhibitor PF-04859989, 20 μL of water was replaced with 20 μL of PF-04859989 at the following concentrations: 100 nM, 1 μM, 10 μM, 100 μM. Following incubation, the tubes were placed on ice, and the reactions were stopped with 50 μL of 6% perchloric acid. The tubes were vortexed and centrifuged (14,000 RPM, 10 minutes), and the supernatant was collected. Samples were stored at −80°C until HPLC analysis for KYNA levels. Protein was evaluated in the homogenate using the Lowry method.^109^

### Experiment 3: Sleep Deprivation (SleepDep) Microdialysis

Sleep deprivation (SleepDep) was conducted during microdialysis in a separate cohort of animals. Male rats received PF-04859989 (30 mg/kg, s.c.) or vehicle (ultrapure water, s.c.) at ZT 23.5 and were subjected to SleepDep from ZT 0 to ZT 6 for a total of 6 hours. Microdialysis chambers were fitted with custom acrylic arms, manually moved by the experimenter to gently sweep across the bottom of the cage and induce wakefulness in a modified gentle handling approach. When the experimenter observed a complete loss of movement or closed eyelids, the bar was manually swept. Microdialysis samples were collected in 30-minute fractions until ZT 6 and in 1-hour fractions from ZT 6 to ZT 12. Food was provided during all microdialysis experiments, though intake was not monitored. Dialysate samples were stored at −80°C until biochemical analyses. Upon completion of the experiment, brains were collected and processed for probe placement, as described above.

### HPLC Analysis of Kynurenic Acid (KYNA)

Twenty μL of microdialysate, diluted 1:2 in ultrapure water, or supernatant collected from the enzyme assays (diluted 1:10 for 2 μM kynurenine assays or 1:100 for 100 μM kynurenine assays in ultrapure water) were analyzed with a Reprosil-Pur C18 column (4 × 150 mm; Dr. MaischGmbh, Ammerbuch, Germany) using a 50 mM sodium acetate mobile phase (pH adjusted to 6.20 with glacial acetic acid) containing 5% acetonitrile at a flow rate of 0.5 mL/minute. KYNA (excitation 344 nm, emission 398 nm) was detected fluorometrically in the eluate (Waters Alliance, 2475 fluorescence detector, Bedford MA) with a post-column addition of 500 mM zinc acetate delivered at a flow rate of 0.1 mL/minute. Data was analyzed using Empower 3 software (Waters). Microdialysis data were not corrected for recovery from the microdialysis probe.

### Statistics

Our studies investigating kynurenine pathway metabolism and extracellular KYNA in rats have noted sex-specific findings.^16, 53, 54^ As such, we designed the present study to analyze the data separately by sex. Data normality was evaluated using the Shapiro–Wilk test and visually inspected with Q-Q plots to confirm an approximately bell-shaped distribution and the absence of outliers. All statistical analyses were performed using GraphPad Prism 10.5.0 software (GraphPad Software, La Jolla, CA, USA) with statistical significance defined as *p* < 0.05.

Supplemental statistical file is included to present all ANOVA results. The results presented in each figure are limited to the significant main effects of time, phase, and/or treatment and a line in each figure legend explains the appropriate annotations.

#### Experiment 1

PF-04859989 microdialysis data (figures 2B/3B) were analyzed separately by phase (light phase beginning at ZT 0 or dark phase beginning at ZT 12) and sex (male or female) using two-way repeated measures (RM) ANOVAs with ZT as a within-subject factor and treatment (vehicle or PF-04859989) as a between-subject factor. Significant main effects were followed up with Fisher’s LSD post hoc tests. Area under the curve analysis (Figures 2C/3C), calculated with data points (as above) relative to Y = 0, was analyzed separately by sex using two-way ANOVAs with phase (light or dark) and treatment (vehicle or PF-04859989) as between-subject factors. For male AUC, any subjects with undetectable KYNA levels by ZT 5 or ZT 17 were not included for AUC analysis (1 light phase and 1 dark phase PF-04859989-treated animals). Significant main effects were followed up with Fisher’s LSD post hoc tests. Kynurenine challenge microdialysis data (Figures 2D/3D) were analyzed for all collected data points separately by phase (light or dark) and sex (male or female), using two-way RM ANOVAs with ZT as a within-subject factor and treatment (vehicle, kynurenine, or PF-04859989 + kynurenine) as a between-subject factor. Significant main effects were followed up with Dunnett’s post hoc tests to vehicle. Area under the curve (Figures 2E/3E), calculated with all measured data points relative to Y=0, was analyzed separately by sex using two-way ANOVAs with phase (light or dark) and treatment (vehicle, kynurenine, or PF-04859989 + kynurenine) as between-subject factors. Significant main effects were followed up with Fisher’s LSD post hoc tests. Female microdialysis data grouped by estrous phase (Supplemental Figure 2) were analyzed with light and dark phase data combined, as no significant phase effects were observed in the female data.

#### Experiment 2

KAT II activity data (Figure 4B/4C) were analyzed by two-way RM ANOVAs with kynurenine concentration as a within-subject factor and ZT as a between-subject factor. Enzyme activity data comparing ZT 3 and ZT 15 (Figure 4D/4E) were analyzed with Mann-Whitney tests at each concentration. PF-04859989 KAT II inhibition data (Figure 4F/4G) were analyzed by two-way RM ANOVAs with PF-04859989 concentration as a within-subject factor and ZT as a between-subject factor. Significant main effects of ZT were followed up with Fisher’s LSD post-hoc tests.

#### Experiment 3

Microdialysis data from SleepDep experiments were calculated as a percentage of basal KYNA and analyzed with a three-way RM ANOVA with ZT as a within-subject factor and SleepDep and PF-04859989 as between-subject factors. Significant main effects of SleepDep or PF-04859989 were followed up with two-way RM ANOVAs separated by SleepDep or PF-04859989 and Fisher’s LSD post-hoc tests compared to control. The area under the microdialysis data curve, calculated using all measured data points relative to Y = 100, was analyzed using a two-way ANOVA with SleepDep and PF-04859989 as between-subject factors, followed by Fisher’s LSD post hoc tests.

##### Safety

This work is not associated with any unexpected, new, or significant hazards or risks.

## Abbreviations

SZ: schizophrenia
BPD: bipolar disorder
α7nAChα7: nicotinic acetylcholine
NMDA: N-methyl-d-aspartate
KYNA: kynurenic acid
SleepDep: sleep deprivation
ZT: Zeitgeber time
KAT: kynurenine aminotransferas
ROS: reactive oxygen species
OAT: organic anion transporter
BBB: blood-brain barrier
P5P: pyridoxal 5’-phosphate

## Author Contributions

**Courtney J. Wright**: Investigation, methodology, formal analysis, visualization, writing – original draft, writing – review and editing. **Silas A. Buck**: Investigation, methodology, formal analysis, writing – review and editing. **Snezana Milosavljevic**: Investigation, formal analysis, writing – review and editing. **Ashley M. Lewis**: Investigation, methodology, formal analysis. **Nathan T.J. Wagner**: Investigation, methodology, formal analysis, visualization, writing – original draft. **Ana Pocivavsek**: Investigation, methodology, formal analysis, visualization, writing – original draft, writing – review and editing, conceptualization, funding acquisition, project administration, supervision, resources.

## Data Availability

The data that support the findings of this study are available from the corresponding author upon reasonable request.

---

**Supplemental Figure 1:**
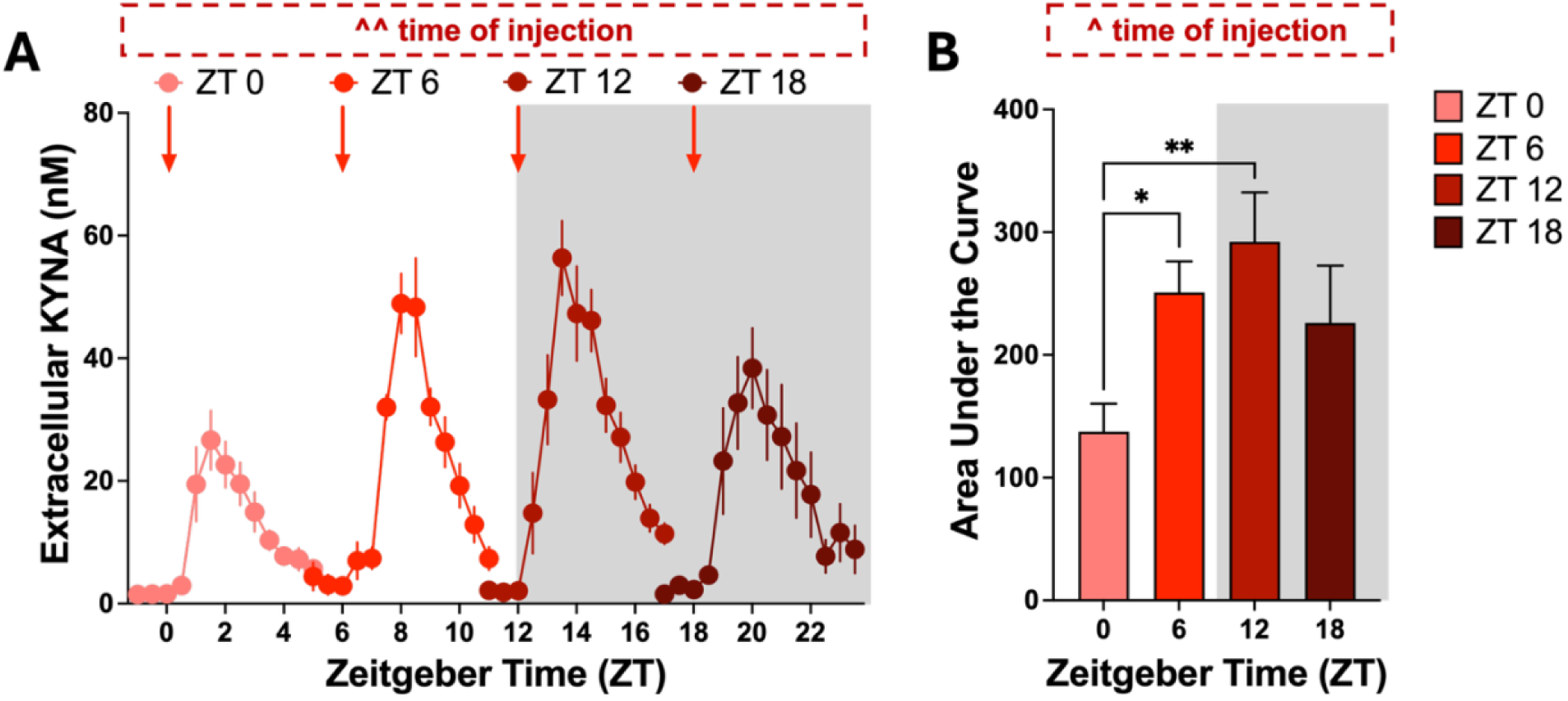
Time of day differentially affects KYNA production following an acute kynurenine challenge, and maximal *de novo* KYNA synthesis occurs at the start of the dark phase. Kynurenine (100 mg/kg, i.p.) was administered at Zeitgeber time (ZT) 0, ZT 6, ZT 12, and ZT 18 in adult male Wistar rats. Extracellular KYNA, collected via in vivo microdialysis, was evaluated in the dorsal hippocampus for the duration of the experiment. **(A)** Extracellular KYNA (two-way RM ANOVA, sample number x time of injection: *F*_9.829, 71.47_ = 3.250, ^^ *p* < 0.01, sample number: *F*_3.273, 71.47_ = 55.20, ^^^^ *p* < 0.0001; time of injection: *F*_3, 23_ = 5.346, ^^ *p* < 0.01). **(B)** Area under the curve (AUC) of extracellular KYNA (one-way ANOVA, *F*_3, 23_ = 4.503, ^ *p* < 0.05). Significant main effects of time (“time of injection”) are displayed in boxed annotations above each figure panel. Data are mean ± SEM and Fisher’s LSD post-hoc test: * *p* < 0.05, ** *p* < 0.01. N = 5 - 10 per group.

**Table 1:**
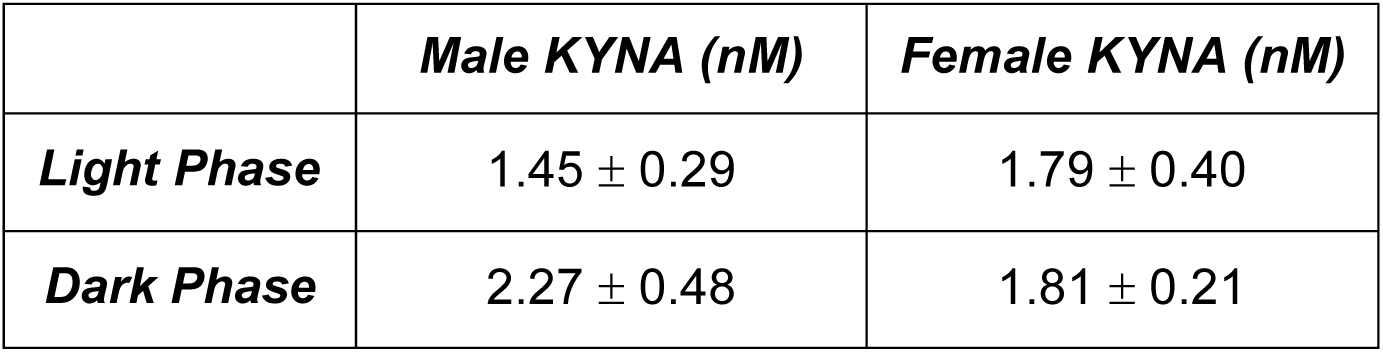
Extracellular KYNA (nM) levels averaged from microdialysis collected during the light or dark phase are not significantly different between male and female rats. Data were calculated as the average KYNA for vehicle-treated rats at zeitgeber time (ZT) 0 – ZT 5 (light phase) or ZT 12 – ZT 17 (dark phase). Two-way ANOVA not significant. Data are expressed as mean ± SEM. N = 8 - 10 per group.

|  | *Male KYNA (nM)* | *Female KYNA (nM)* |
| --- | --- | --- |
| *Light Phase* | 1.45 ± 0.29 | 1.79 ± 0.40 |
| *Dark Phase* | 2.27 ± 0.48 | 1.81 ± 0.21 |

**Supplemental Figure 2:**
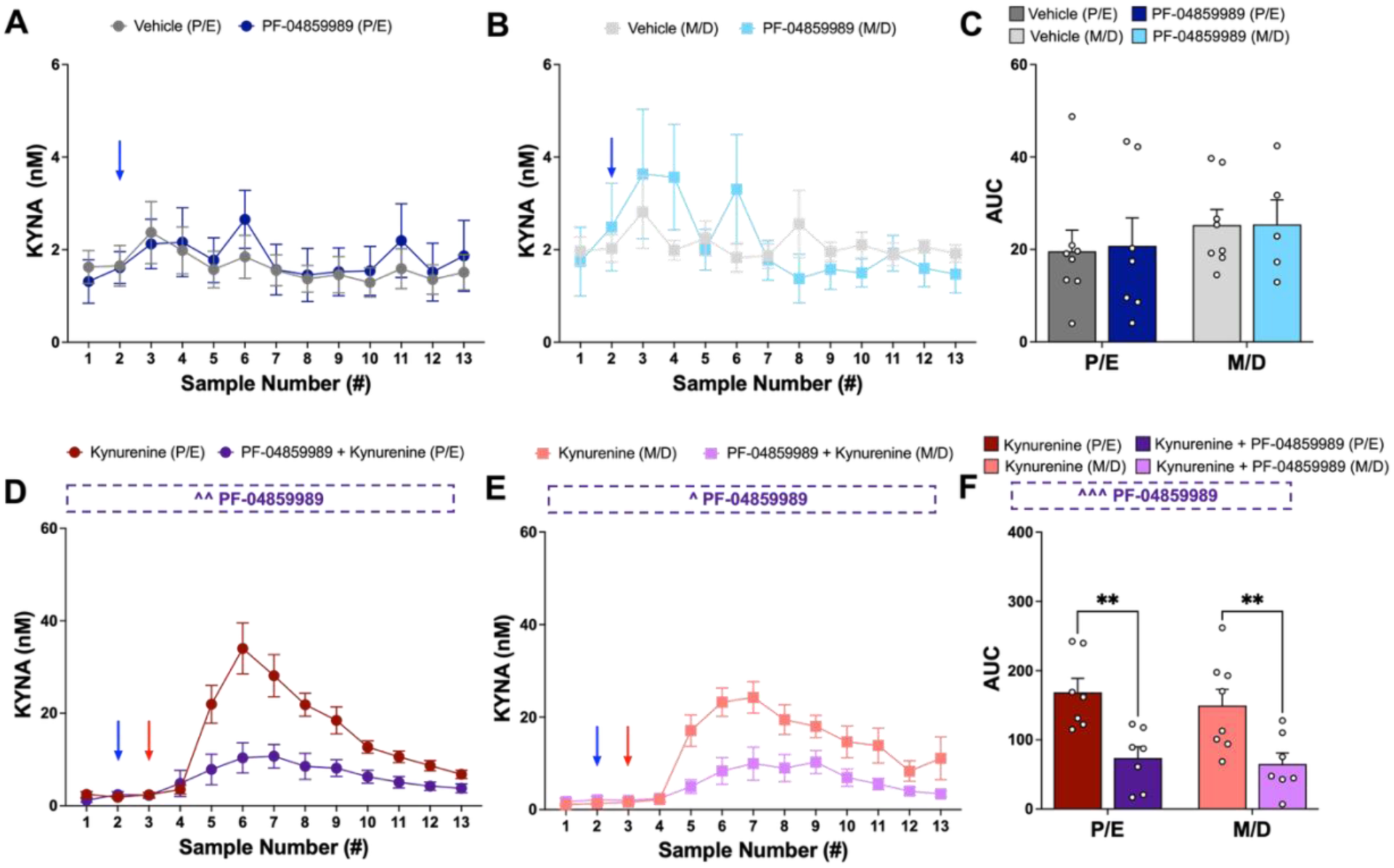
Estrous cycle stage does not impact basal KYNA, response to kynurenine challenge, or PF-04859989 treatment in females. **(A)** Extracellular KYNA in females (light and dark phase data combined) was evaluated in females with high circulating estrogen (**proestrus/estrus, P/E**) upon vehicle or PF-04859989 treatment alone (two-way RM ANOVA, ZT: *F*_4.957, 62.79_ = 2.497, ^ *p* < 0.05). **(B)** Extracellular KYNA levels in females with low circulating estrogen (**metestrus/diestrus, M/D**) treated with vehicle or PF-04859989 alone (two-way RM ANOVA, n.s.). **(C)** AUC of extracellular KYNA in females treated with vehicle or PF-04859989 alone, separated by P/E or M/D (two-way ANOVA, n.s.). **(D)** Extracellular KYNA in P/E females upon kynurenine or PF-04859989 + kynurenine treatment (two-way RM ANOVA, treatment x ZT: *F*_2.131, 24.50_ = 8.020, ^^ *p* < 0.01, treatment: *F*_1, 13_ = 13.42, ^^ *p* < 0.01, ZT: *F*_2.131, 24.50_ = 25.25, ^^^^ *p* < 0.0001). **(E)** Extracellular KYNA in M/D females (light and dark phase data combined) upon kynurenine or PF-04859989 + kynurenine treatment (two-way ANOVA, treatment x ZT: *F*_2.848, 35.37_ = 4.809, ^^ *p* = 0.0073, ZT: *F*_2.848, 35.37_ = 18.90, ^^^^ *p* < 0.0001, treatment: F_1,13_ = 6727, ^ *p* = 0.0223). **(F)** Area under the curve (AUC) of extracellular KYNA in females treated with kynurenine or PF-04859989 + kynurenine, separated by P/E or M/D (two-way ANOVA, treatment: *F*_1, 25_ = 21.11, ^^^ *p* = 0.0001). Significant main effects of PF-04859989 are displayed in boxed annotations above each figure panel. All data are mean ± SEM. Fisher’s LSD post-hoc: \**p* < 0.05, \*\**p* < 0.01. N = 5 - 8 per group.

**Supplemental Figure 3:**
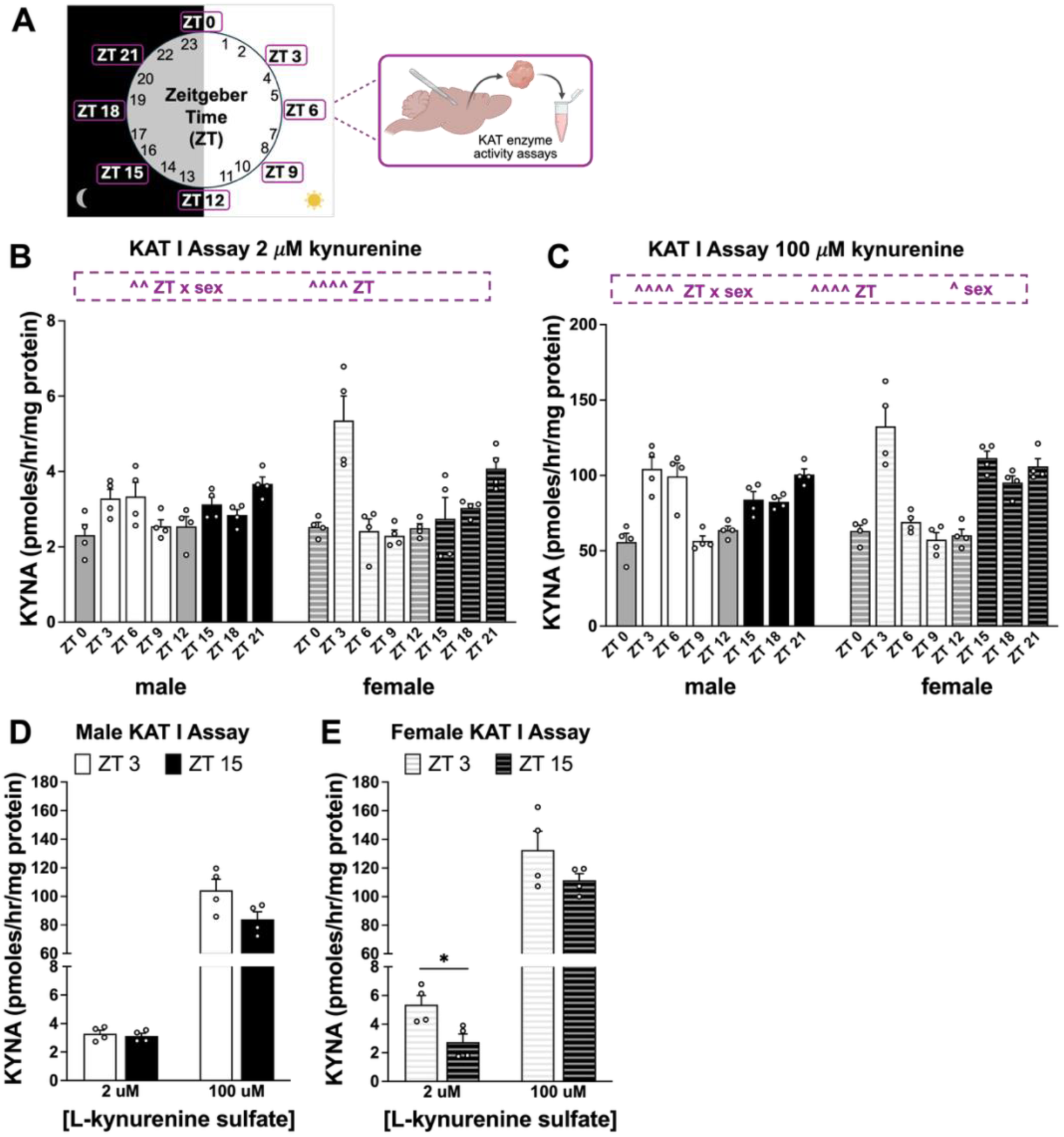
Kynurenine aminotransferase I (KAT I) enzyme activity changes across Zeitgeber time (ZT). **(A)** Experimental approach. Brain samples were collected from male and female rats at 3-hour intervals for *ex vivo* enzyme activity assays, KAT I and KAT II, which were assessed by measuring kynurenic acid (KYNA) production. **(B)** Forebrain KAT I enzyme activity assay, 2 *μ*M kynurenine (two-way ANOVA, ZT x sex: *F*_7, 48_ = 4.252, *p* < 0.01; ZT: *F*_7, 48_ = 10.79, *p* < 0.0001). **(C)** Forebrain KAT I enzyme activity assay, 100 *μ*M kynurenine (two-way ANOVA, ZT x sex: *F*_7, 48_ = 5.199, *p* < 0.001, ZT: *F*_7, 48_ = 30.68, *p* < 0.0001, sex: *F*_1, 48_ = 4.337, *p* < 0.05). **(D)** Male KAT I activity at ZT 3 and ZT 15 (Mann-Whitney tests n.s.). **(E)** Female KAT I activity at ZT 3 and ZT 15 (Mann-Whitney test ZT 3: \**p* < 0.05, ZT 15: n.s.). Significant interactions and main effects are displayed in boxed annotations above each figure panel. All data are mean ± SEM. N = 4 per group.

**Supplemental Figure 4:**
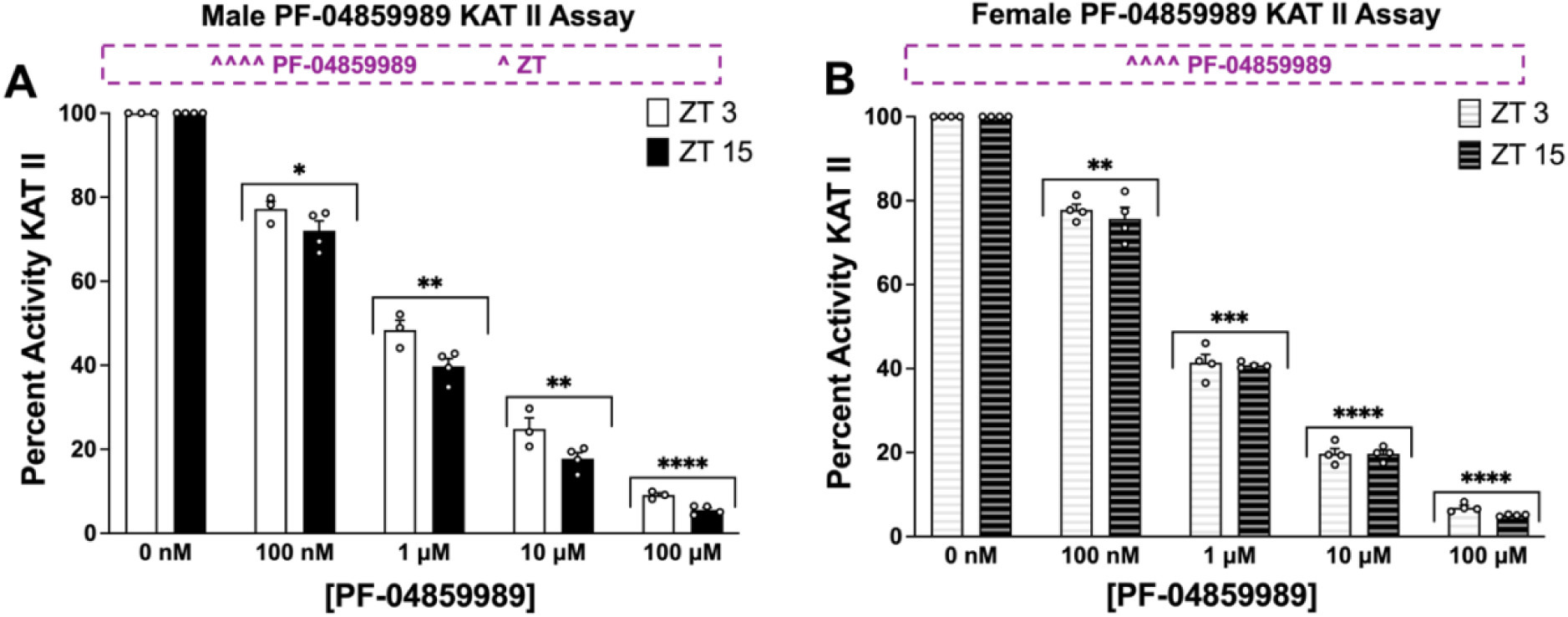
PF-04859989 significantly inhibits kynurenine aminotransferase (KAT) II enzyme activity in a dose-dependent manner. **(A)** Male forebrain percent activity KAT II with increasing concentrations of PF-04859989 (two-way ANOVA, PF-04859989 concentration: *F*_2.065, 10.33_ = 1516, ^^^^ *p* < 0.0001, ZT: *F*_1, 5_ = 10.62, ^ *p* < 0.05). **(B)** Female forebrain percent activity KAT II with increasing concentrations of PF-04859989 (two-way ANOVA, PF-04859989 concentration: *F*_1.978, 11.87_ = 2269, ^^^^ *p* < 0.0001). Significant interactions and main effects are displayed in boxed annotations above each figure panel. All data are mean ± SEM and Dunnett’s post-hoc comparison to 0 nM PF-04859989. Post-hoc significance denotes the least significant ZT 3 or ZT 15 comparison between 0 nM and subsequent PF-04859989 concentrations: \**p* < 0.05, \*\**p* < 0.01, \*\*\**p* < 0.001, \*\*\*\**p* < 0.0001. N = 3 – 4 per group.

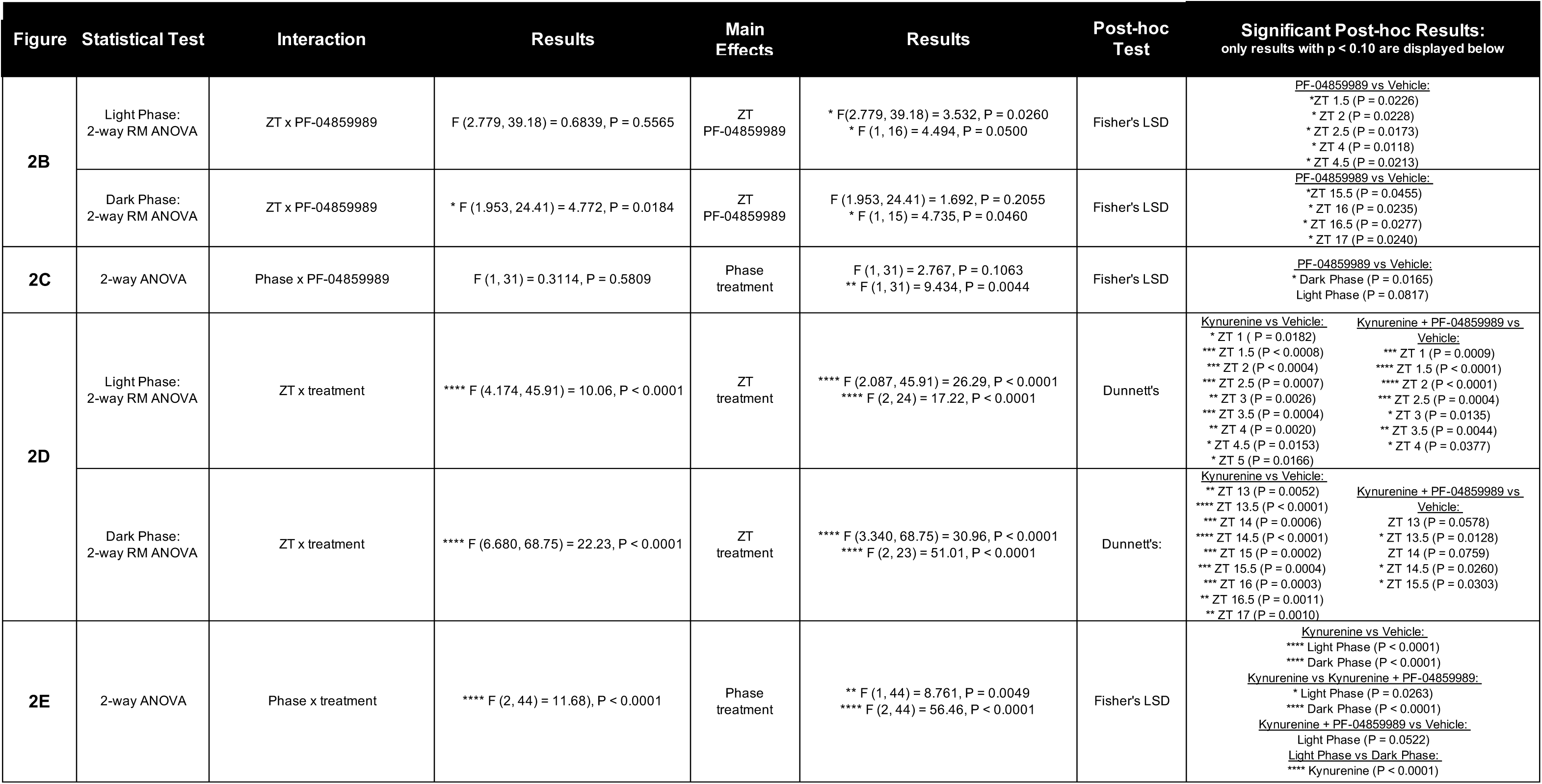
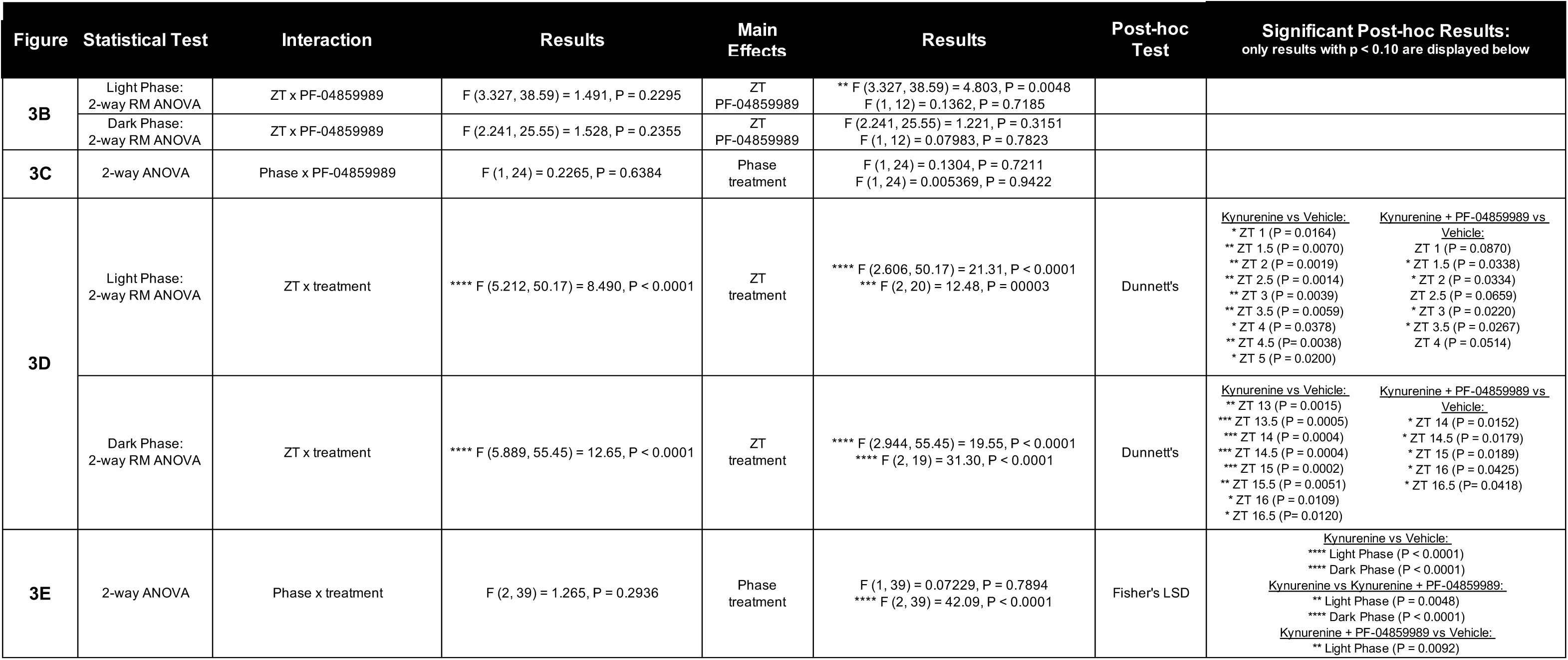
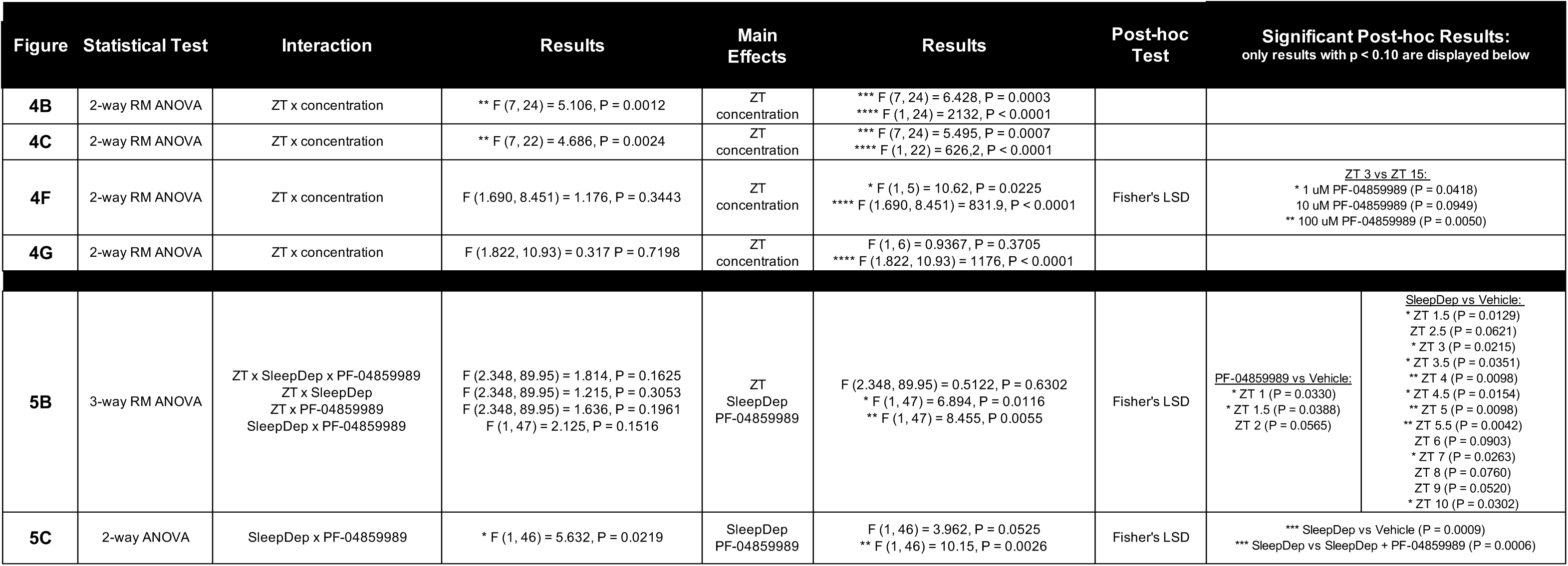
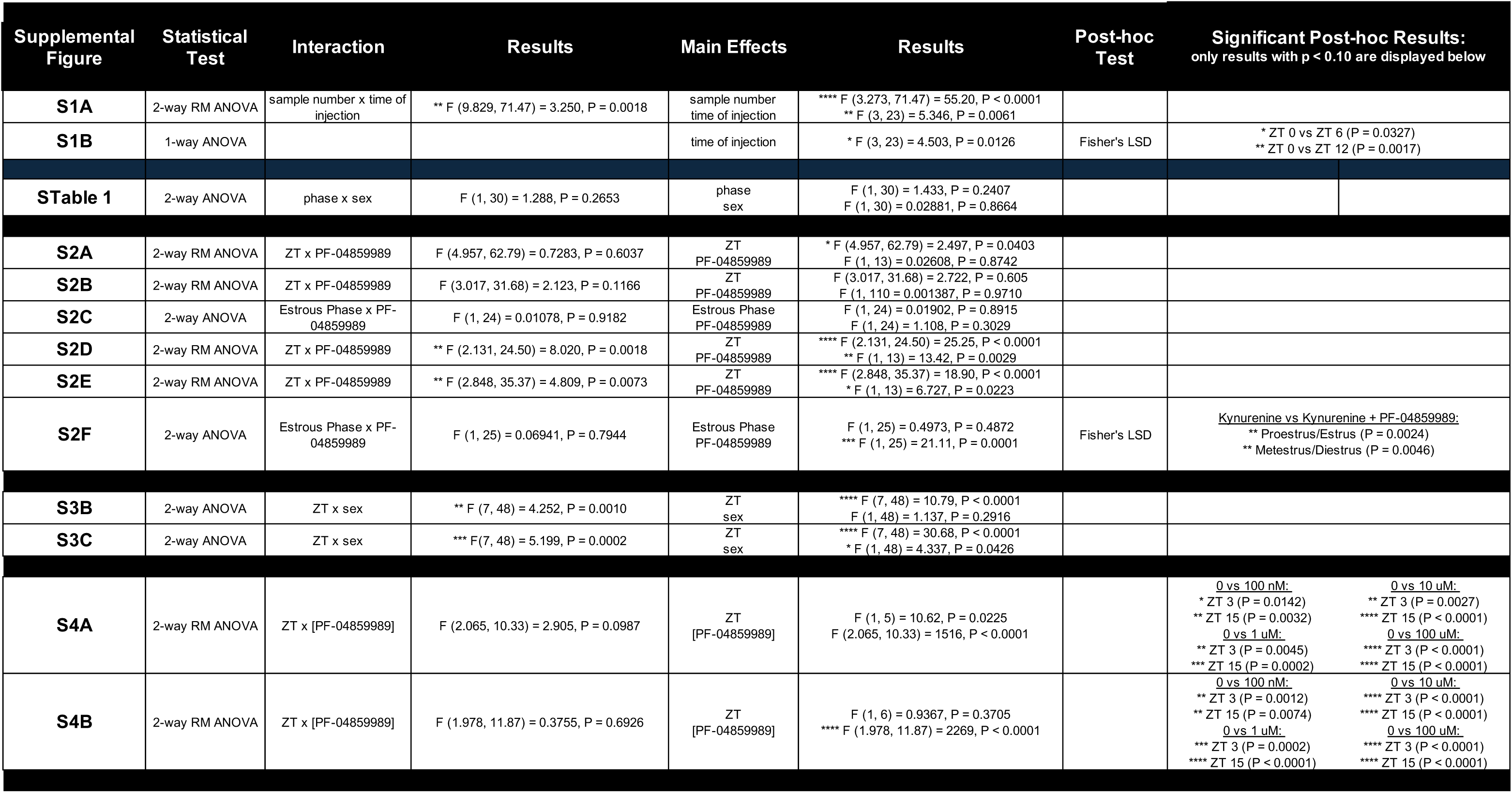

